## Supplemental Materials for "HeALTH: An Automated Platform for Long-term Longitudinal Studies of Whole Organisms under Precise Environmental Control"

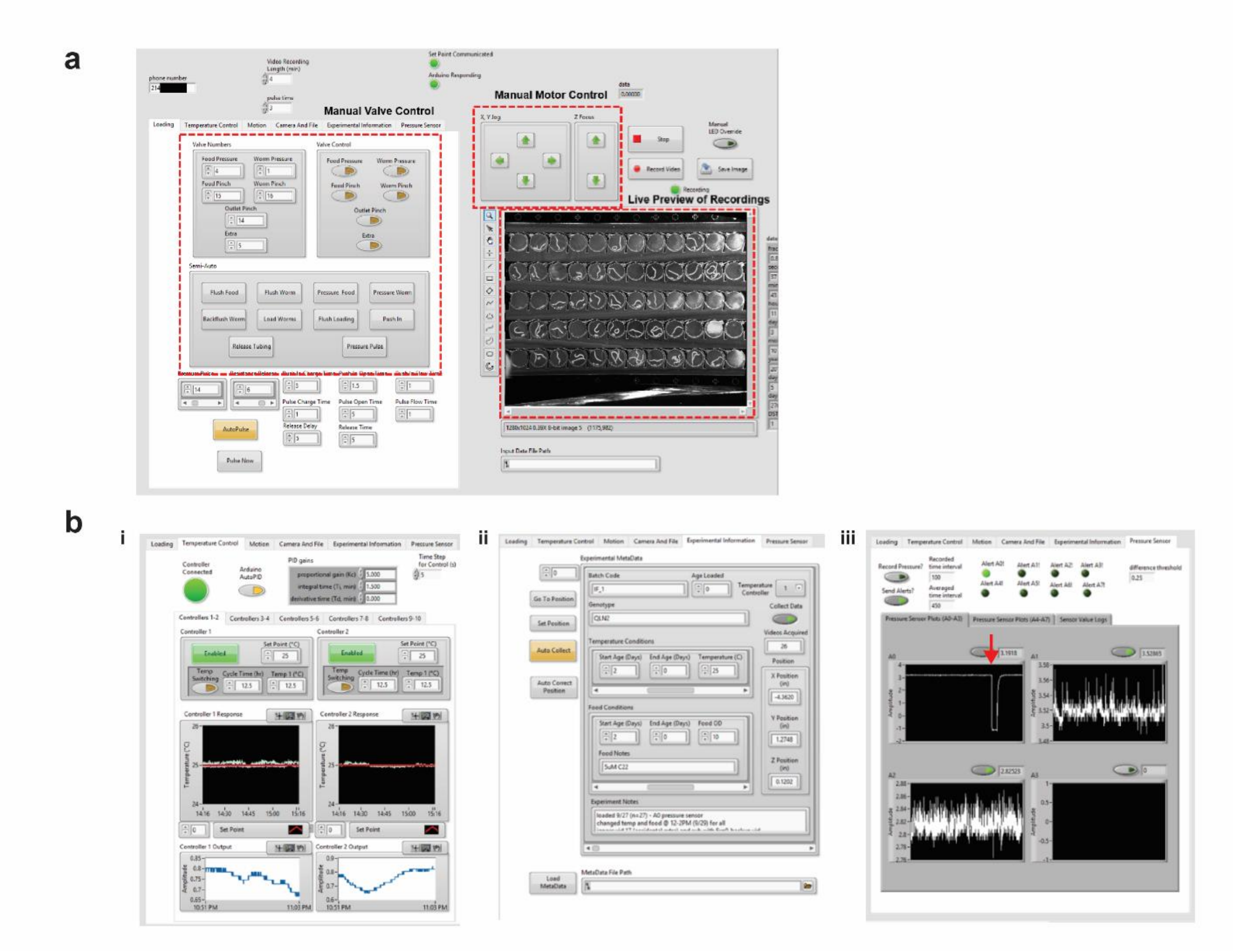

**Supplementary Figure 1.** HeALTH experimental GUI

a) Main page of the user GUI. This allows experimenters to manually control valves for fluid flow, move the camera and/or stage, and view/take records of devices in real-time. b) Additional subpanels of the GUI allowing for additional user control and input. i) Individual temperature control and monitoring for each detected temperature module. ii) Metadata and experimental information tracking for each device. iii) Continuous pressure monitoring and readout for automated clog detection. The drop in pressure (red arrow) indicates the opening of a downstream valve.

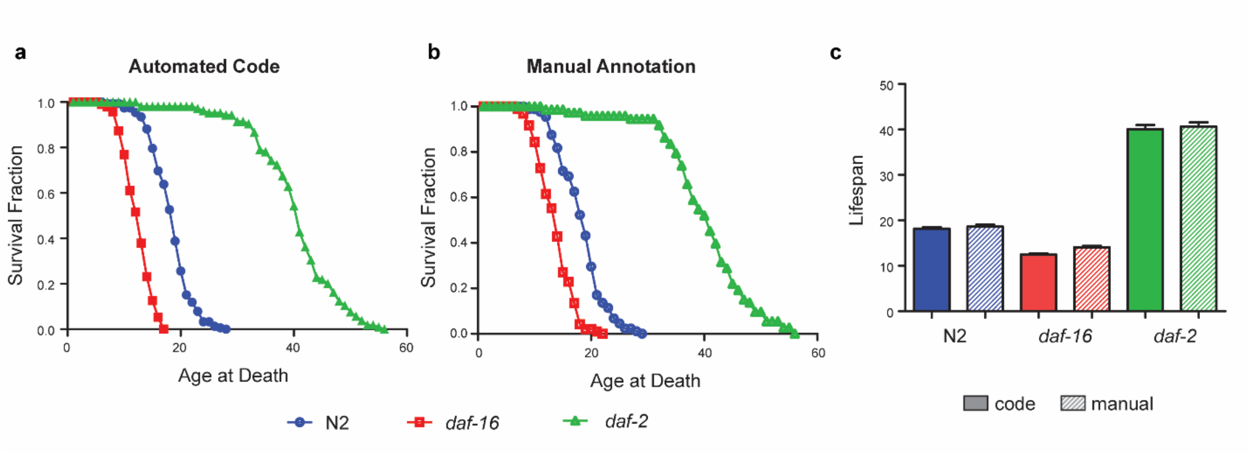

**Supplementary Figure 2.** Validation of automated live/dead code

a) Lifespan curves obtained from the automated analysis code for wild-type (18.20 days ± 0.37, n = 87 individuals), *daf-16* (12.48 days ± 0.26, n = 95 individuals), and *daf-2* (40.08 days ± 0.91, n = 73 individuals) populations. Error is reported as SEM. b) Manual annotation of death for the videos inputted into the automated analysis code for wild-type (18.48 days ± 0.42), *daf-16 (*13.78 days ± 0.32), and *daf-2* (40.29 days ± 0.94) populations. Error is reported as SEM. c) Comparison of average lifespans for wild-type, *daf-16*, and *daf-2* populations as found by the code and manual annotation. Error is reported as SEM. No significant difference was found between code outputted and manually annotated lifespans (log-rank test, *daf-16* p = 0.0162, N2 p = 0.3609, *daf-2* p = 0.6333)

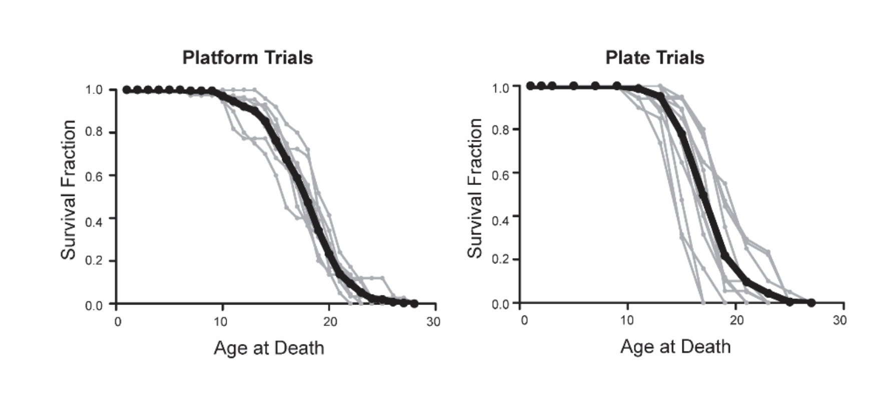

**Supplementary Figure 3.** Characterization of trial-to-trial variability

a) Lifespan curves for each trial for N2, cultured at 25°C with $OD_{600}5$ food concentration on the platform. Gray curves show individual trials while the black curve shows the aggregated lifespan curve. b) Lifespan curves for each trial for N2, cultured at 25°C with $OD_{600}5$ food concentration on plate. Gray curves show individual trials while the black curve shows the aggregated lifespan curve.

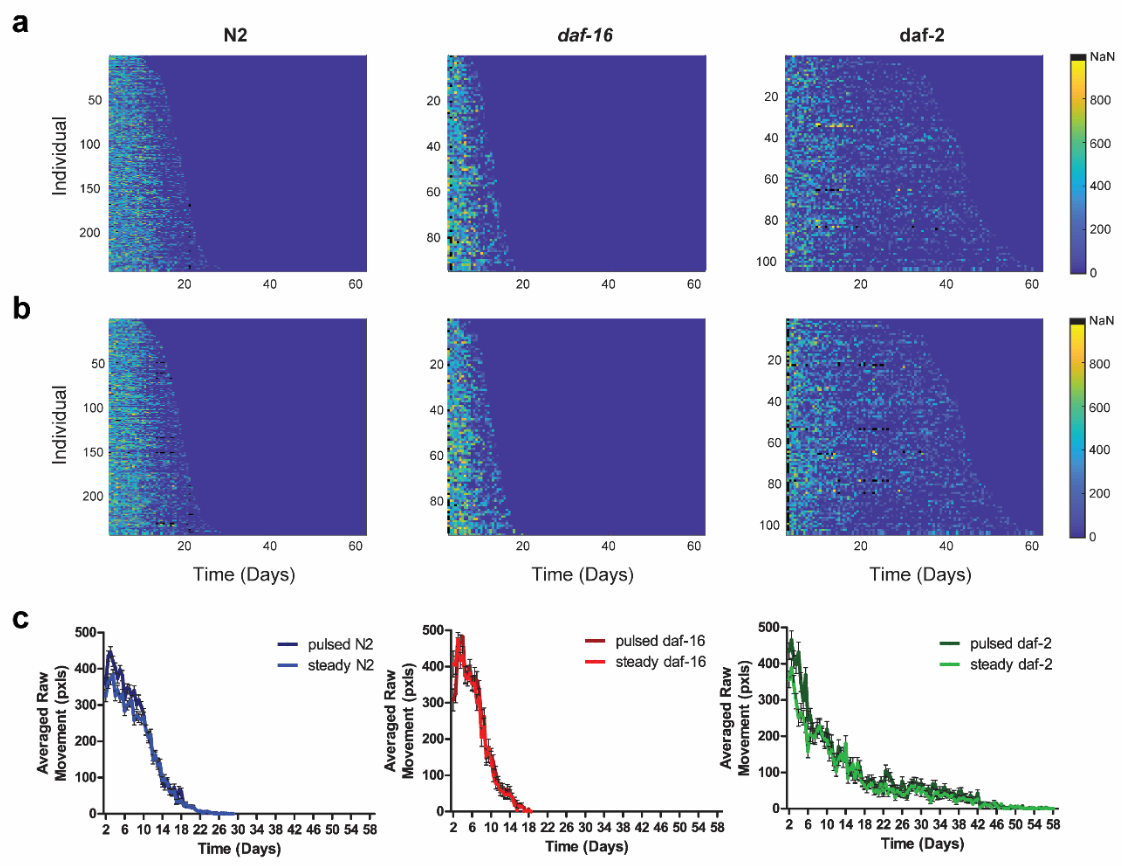

**Supplementary Figure 4.** Worms under constant fluid-flow demonstrated evoked behavior

a) Heatmaps of individual raw movement over time for N2, *daf-16,* and *daf-2* populations under constant flow conditions (~15µL/min). b) Heatmaps of raw movement over time for N2, *daf-16,* and *daf-2* populations after opening the downstream solenoid valve, creating a strong, pulse of fluid flow (~275µL/min) for 5 seconds. c) Population averages of raw movement over time for both the constant flow and 'pulsed' flow conditions across genotypes. No statistical significance was found between the two flow conditions (Kolmogorov-Smirnov test, N2 p = 0.9782, *daf-16* p = 1.000, *daf-2* p = 0.1413), indicating the constant flow conditions create enough of a mechanical stimulus to view stimulated behavior. Error bars are SEM.

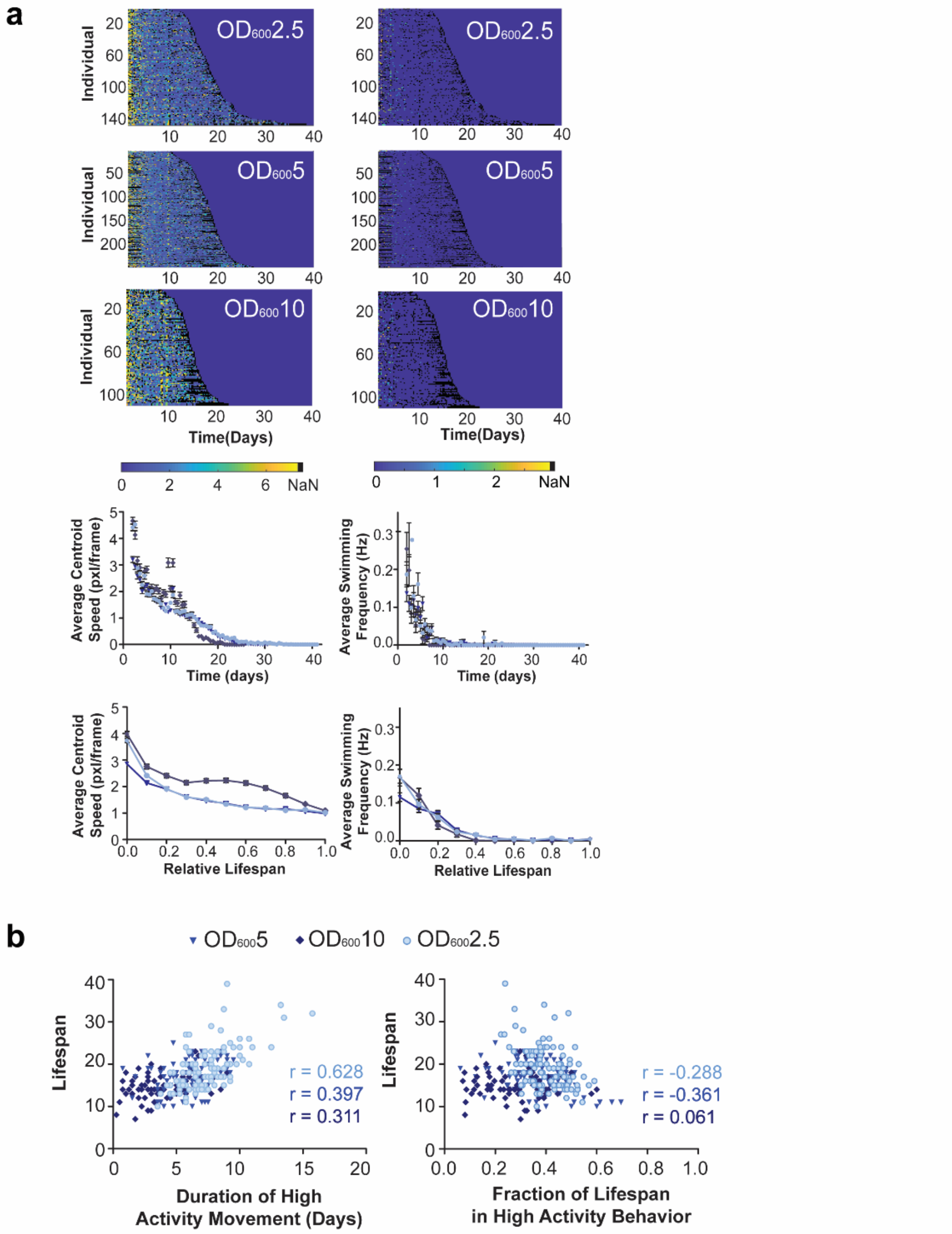

**Supplementary Figure 5.** Extended longevity and behavioral analysis for different food levels

a) (*left)* Heatmaps of individual maximum centroid speed, population averaged activity, and relative population average normalized to lifespan across different food levels. Error bars are SEM. (*right)* Heatmaps of individual swimming frequency, population averaged activity, and relative population average normalized to lifespan across different food levels. Error bars are SEM. b) Scatter plots of duration of high activity for raw movement vs. individual lifespan and fraction of life with high activity for raw movement vs. individual lifespan across food levels. Pearson correlation coefficients are listed for each food level.

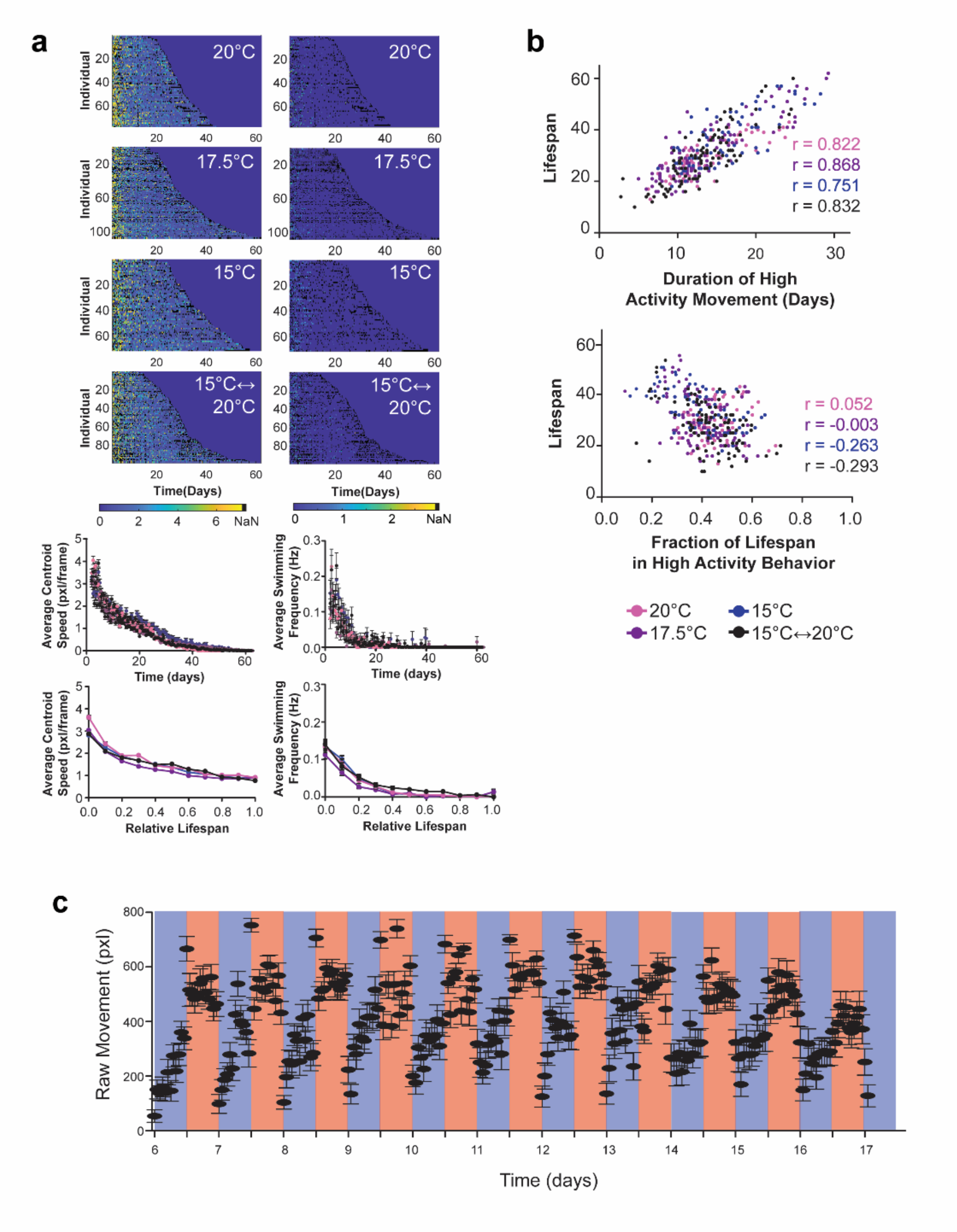

**Supplementary Figure 6.** Extended longevity and behavioral analysis for thermal perturbations

a) (*left)* Heatmaps of individual maximum centroid speed, population averaged activity, and relative population average normalized to lifespan across different temperature conditions. Error bars are SEM. (*right)* Heatmaps of individual swimming frequency, population averaged activity, and relative population average normalized to lifespan across different temperature conditions. Error bars are SEM. b) Scatter plots of duration of high activity for raw movement vs. individual lifespan and fraction of life with high activity for raw movement vs. individual lifespan across temperature conditions. Pearson correlation coefficients are listed for each thermal condition. c) Average raw movement of a population under oscillatory thermal swings (n = 26 individuals). Error bars are SEM. Blue background indicates a culture temperature of 15°C and a red background indicates a temperature of 20°C.

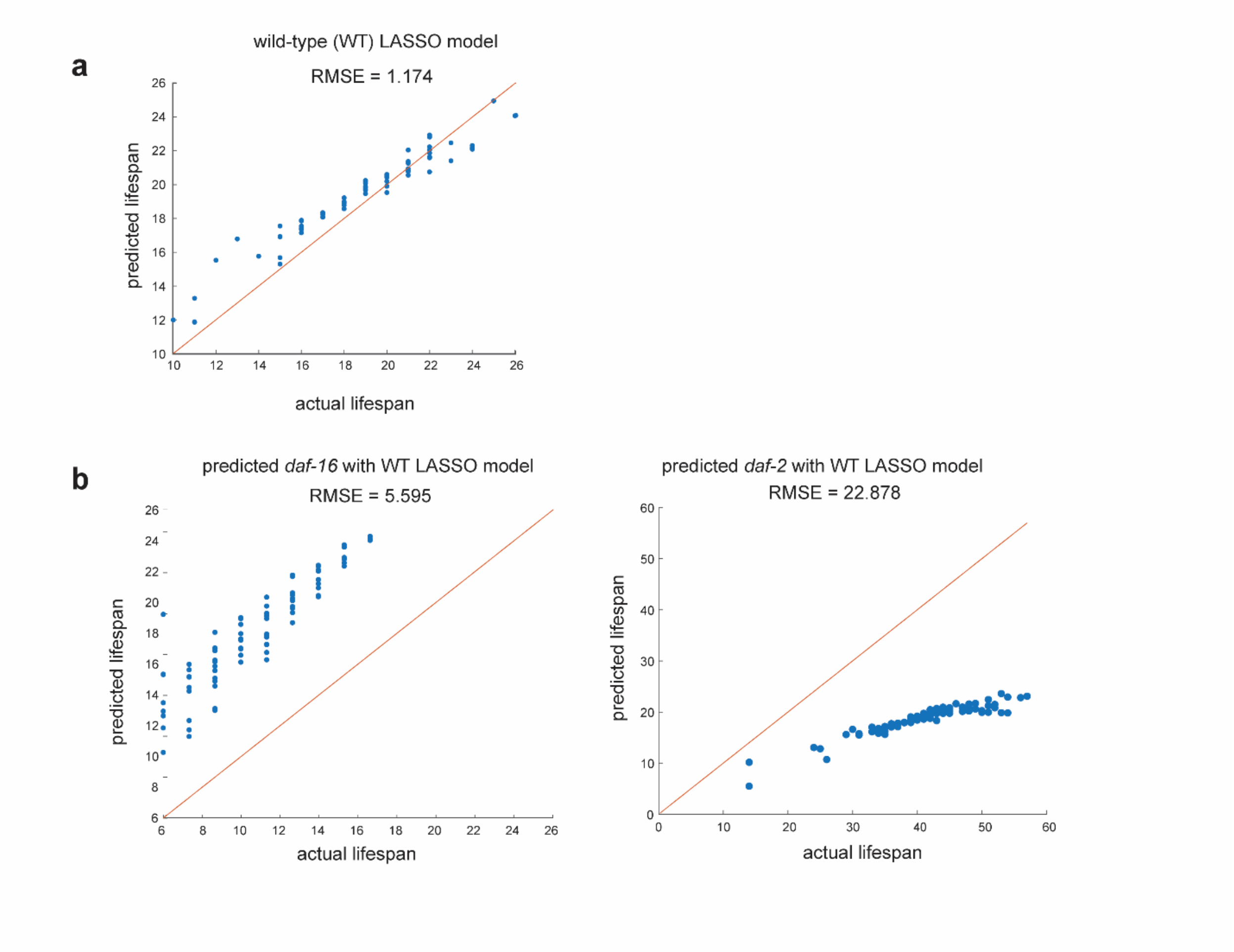

**Supplementary Figure 7.** Predictive models of lifespan based on behavioral decline

a) Comparing predicted lifespan v. actual lifespan across a wild-type population (n = 73 individuals) cultured at 25°C with $OD_{600}$5 using a LASSO regression model. Diagonal line indicates a theoretically perfect prediction. b) Comparing predicted lifespan v. actual lifespan for *daf-16* (n = 95 individuals) and *daf-2* populations (n = 105 individuals) using the corresponding wild-type predictive model (cultured at 25°C with $OD_{600}$5). The model consistently overpredicts the lifespan of *daf-16* individuals, while underpredicting the lifespan of *daf-2* individuals.

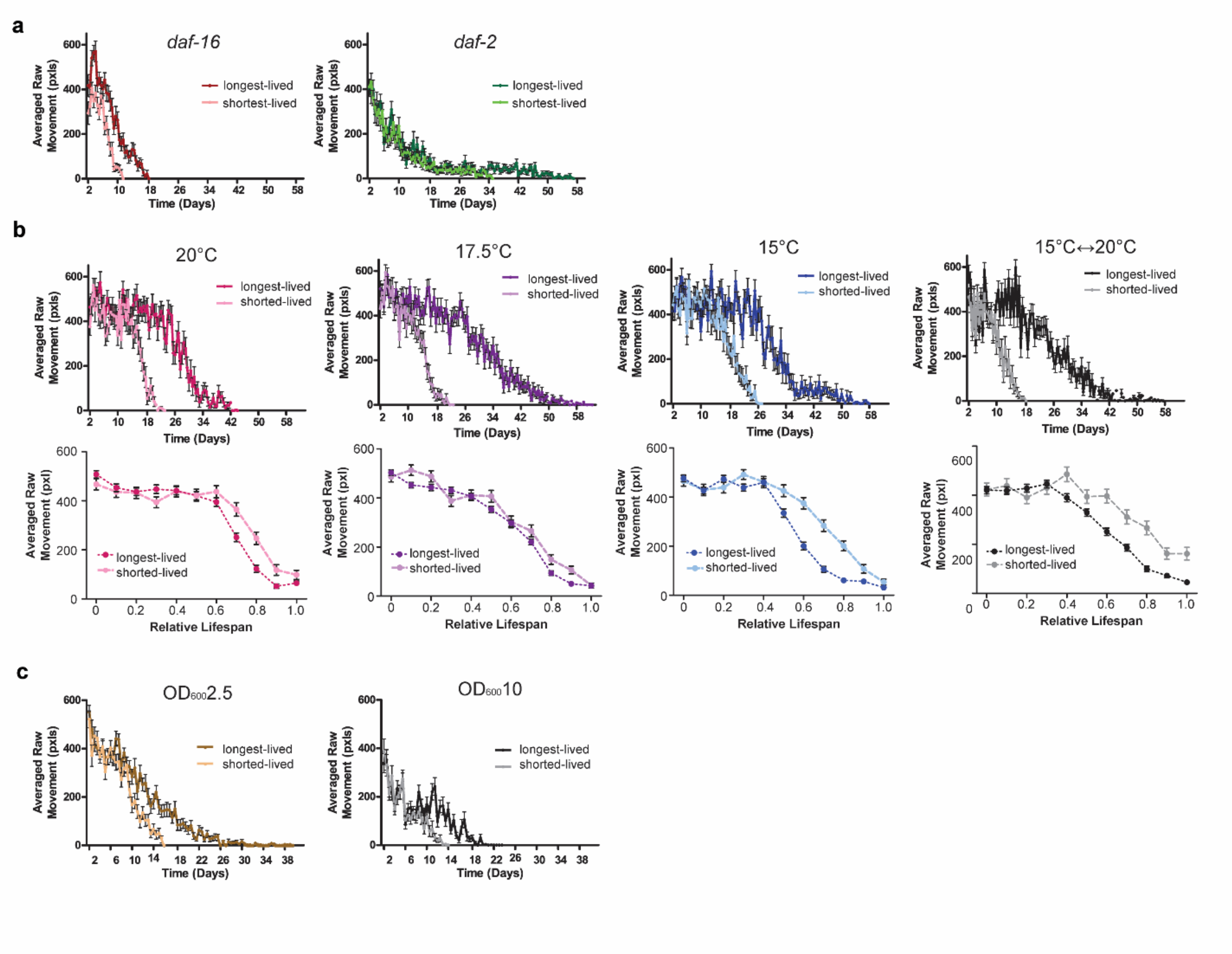

**Supplementary Figure 8.** Intrapopulation behavioral decline across genetic and environmental perturbations

a) Averaged raw movement over time of the short- and long-lived subpopulations at 25°C at $OD_{600}5$ food level across different genotypes. Error bars are SEM. b) (*top)* Averaged raw movement over time of the short- and long-lived wild-type subpopulations at $OD_{600}5$ food level across different thermal conditions. 20°C short- (n = 16 individuals) and long-lived (n = 16 individuals) subpopulations. 17.5°C short- (n = 24 individuals) and long-lived (n = 22 individuals) subpopulations. 15°C short- (n = 14 individuals) and long-lived (n = 15 individuals) subpopulations. 15°C<->20°C short- (n = 21 individuals) and long-lived (n = 22 individuals) subpopulations. Error bars are SEM. *(bottom)* Corresponding average raw movement over normalized, relative lifespan of the short- and long-lived subpopulations across different thermal conditions. Error bars are SEM. c) Averaged raw movement over time of the short- and long-lived wild-type subpopulations at 25°C across different food levels. Error bars are SEM.

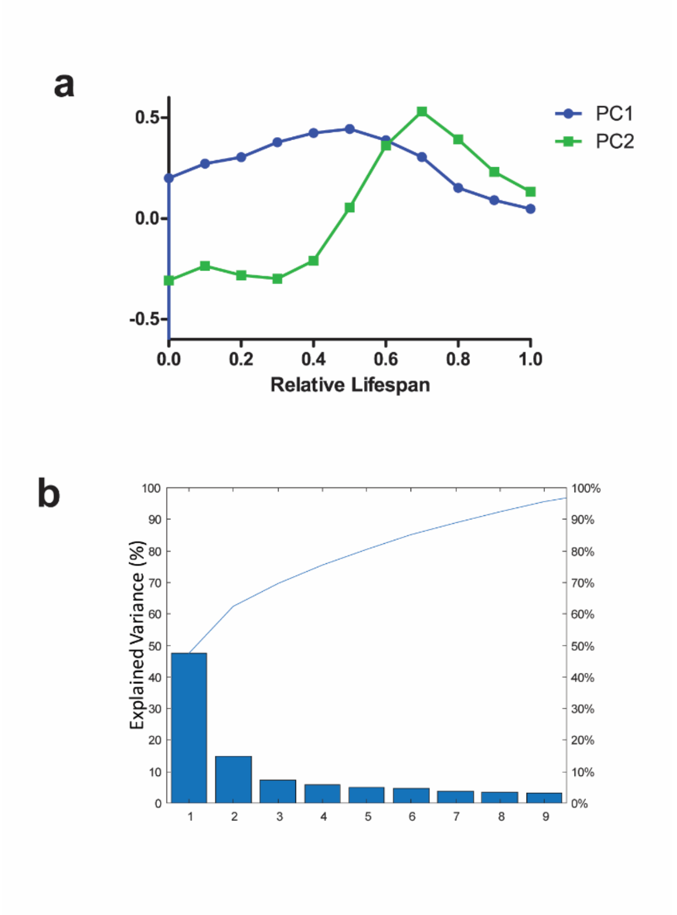

**Supplementary Figure 9.** Characterization of PCA

a) First two principle component curves. b) Scree plot of explained variance with each principle component.

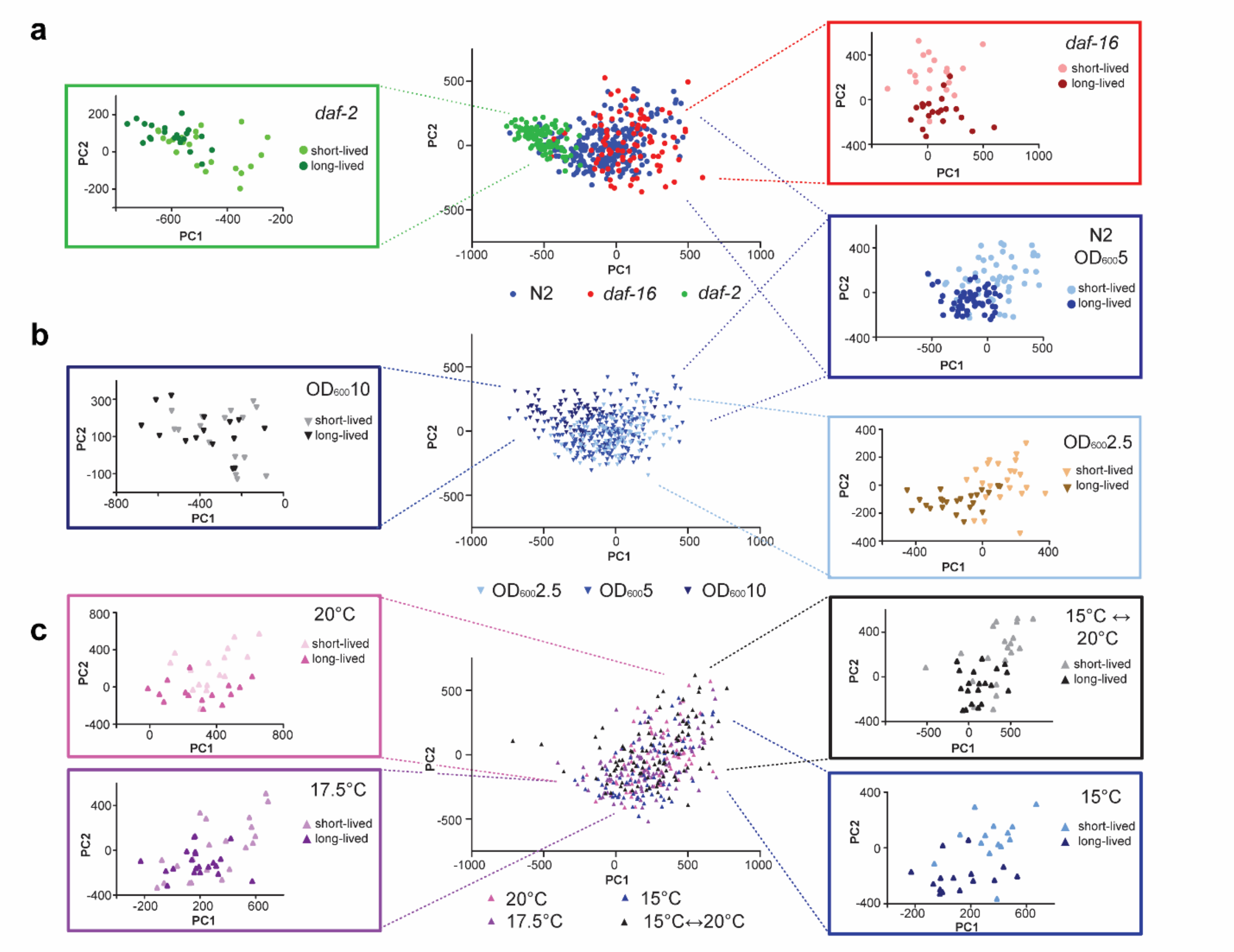

**Supplementary Figure 10.** Intrapopulation variation within the PC space

a) Individuals cultured at 25°C at food level $OD_{600}5$. Inserts show the spatial location and division of the short-lived and long-lived cohorts across the different genotypes. b) Wild-type individuals cultured at 25°C. Inserts show the spatial location and division of the short-lived and long-lived cohorts across the different food levels. c) Wild-type individuals cultured at food level $OD_{600}5$. Inserts show the spatial location and division of the short-lived and long-lived cohorts cultured at different thermal conditions.

**Supplementary Videos 1-3.** Time-lapse videos of a wild-type worm in device cultured at 25°C at $OD_{600}5$

Compiled time-lapse video of an individual cultured on HeALTH from the initial L4 stage to the end of the lifespan. The wild-type (N2) worm is cultured at 25°C at $OD_{600}5$ from Day 2 adult to death. From L4 larval stage to Day 1 adult the worm was cultured in $OD_{600}10$ and at 20°C to ensure no adverse developmental effects from low food levels or increased temperature. The video is shown at twice the speed.

**Supplementary Videos 4-6.** Time-lapse videos of a wild-type worm in device cultured at 25°C at $OD_{600}10$

Compiled time-lapse video an individual cultured on HeALTH from the initial L4 stage to the end of the lifespan. The wild-type (N2) worm is cultured at 25°C at $OD_{600}10$ from Day 2 adult to death. From L4 larval stage to Day 1 adult the worm was cultured in $OD_{600}10$ and at 20°C to ensure no adverse developmental effects from low food levels or increased temperature. The video is shown at twice the speed.

**Supplementary Videos 7-9.** Time-lapse videos of a wild-type worm in device cultured at 25°C at $OD_{600}2.5$

Compiled time-lapse video an individual cultured on HeALTH from the initial L4 stage to the end of the lifespan. The wild-type (N2) worm is cultured at 25°C at $OD_{600}2.5$ from Day 2 adult to death. From L4 larval stage to Day 1 adult the worm was cultured in $OD_{600}10$ and at 20°C to ensure no adverse developmental effects from low food levels or increased temperature. The video is shown at twice the speed.

**Supplementary Videos 10-12.** Time-lapse videos of a *daf-16* worm in device cultured at 25°C at $OD_{600}5$

Compiled time-lapse video of an individual cultured on HeALTH from the initial L4 stage to the end of the lifespan. The *daf-16* worm is cultured at 25°C at $OD_{600}5$ from Day 2 adult to death. From L4 larval stage to Day 1 adult the worm was cultured in $OD_{600}10$ and at 20°C to ensure no adverse developmental effects from low food levels or increased temperature. The video is shown at twice the speed.

**Supplementary Videos 13-15.** Time-lapse videos of a *daf-2* worm in device cultured at 25°C at $OD_{600}5$

Compiled time-lapse video of an individual cultured on HeALTH from the initial L4 stage to the end of the lifespan. The *daf-2* worm is cultured at 25°C at $OD_{600}5$ from Day 2 adult to death. From L4 larval stage to Day 1 adult the worm was cultured in $OD_{600}10$ and at 20°C to ensure no adverse developmental effects from low food levels or increased temperature. The video is shown at twice the speed.

| | PLATFORM VARIABILITY | | PLATE CONTROL VARIABILITY | |
|---|---|---|---|---|
| Source of Variability | Variance | Percent of Variance (%) | Variance | Percent of Variance (%) |
| Trial | 0.8281 | 6.41 | 2.2211 | 27.16 |
| Trial x Plate | 0.1568 | 1.21 | 0.0237 | 0.29 |
| Individual Variation | 11.9275 | 92.26 | 5.9189 | 72.38 |
| Total | **12.9124** | | **8.1773** | |

**Supplementary Table 1.** Sources of variability across experimental trials

Variance components and the percent of variance due to difference sources of variability across trials cultured on the platform and on plate. Variance was calculated using a GLM mixed-effect model [26]. Total variation across the two culture methods is comparable. Variance attributed to different trials is lower within the platform. The individual variation is higher; however, this could be due to bacteria. A single trial of the plate assay was seeded with the same batch of bacteria, while multiple batches were grown and used on the platform due to the high consumption rate of bacteria culture. As a result, the high 'individual variation' could be due to variation across bacterial cultures.

**Supplementary Note 1.** HeALTH Design Documentation

We present HeALTH, a platform for performing automated longitudinal healthspan studies for *C. elegans* under precise environmental control. The following documentation is a brief overview on how the entire platform was built and customized for the set of experiments shown in the article. By design our system is modular and scalable. As a result, it can be customized depending on the requirements of the user. The complete bill of materials (BOM) is organized by subsystem and is listed at the end of the documentation. Details for the specific parts mentioned in the text can be found in the BOM and are referenced by item number within each system sub-section. All CAD files are uploaded to GrabCAD at: https://workbench.grabcad.com/workbench/projects/gcwDW30oR56MuBI2OsZ-ge4yfAhwFzatnB6-7xhbRppU0R#/space/gcSlzhGMjRykJLl3yZbV99GF6oXy3TtZqJsoVO-CstAUnH

### 1. Imaging System

The imaging system consists of a single camera mounted on an automated x-y stage that travels and record the behavior of populations across different devices on the platform. To create a motorized stage system with a wide range of motion and precise X, Y, and Z directional movement in a simple, efficient manner we modified a commercially available DIY CNC stage kit, where three separate stepper motors driven by a microcontroller are coupled to threaded rods drive motion across the three directions. We chose the MyDIYCNC Desktop CNC Machine from MyDIYCNC. Unfortunately, the product and company now no longer exist; however, the general idea and modifications we made should be translatable to other, currently available DIY stage kits. As a result, the exact dimensions and modifications should be modified as needed to fit the user's actual system. For reference, our system has an X range of motion of ~15", Y motion of ~13", and Z motion of ~3", with a resolution of 10 µm across all directions. The following sections detail how to modify the motorized stage and how to create the imaging and illumination system.

#### 1.1 Modifying the motorized stage

One of the major modifications necessary for many CNC stage kits is to alter the height of the machine in order to allow for the imaging components. To do so, we added height extension panels (#18) to the sides of the assembled CNC stage. Additionally, to make the system more compatible with fluids, we altered the stage material from patterned metal sheet to acrylic (15"L x 13"W x 3/16", #19). We also removed the router mount and replaced it with a custom machined HDPE part that holds and connects the lens clamp (#4). Based on the type of kit purchased, the screw holes will need to be adjusted; however, the holder for the clamp should be constant. See below for an image of the imaging system.

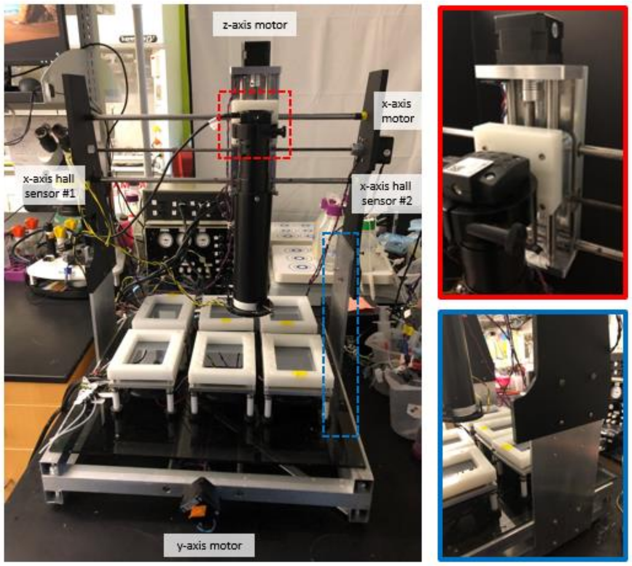

**Overview of the imaging system**. (*left*) Image of imaging system with temperature modules on the stage. Major components of the motorized stage are labeled. (*top right, red insert*) close-up of the machined HDPE lens clamp holder. *(bottom right*, blue insert) close-up of the machined height extension panels

To more easily integrate motor control with the other platform components we rerouted the motor wiring (#16-17) to a small microcontroller (#23, #26-28), which then communicates with the integrated LabVIEW GUI. The microcontroller outputs digital pulses to drive motion from the stepper motors. We also added hall effect proximity sensors (#20) on the edges of each axis for positional information and connected it to the microcontroller as a limit switch control. The x-axis halls sensor locations are labeled in the image shown above. The motor circuit is housed in a custom machined project box (6.112" L x 4.612" W X 2.356", #22). The system is powered by a 12V power supply (#24-25). See below for the electronic circuit diagram, illustrating the connections for a single motor and its corresponding sensors.

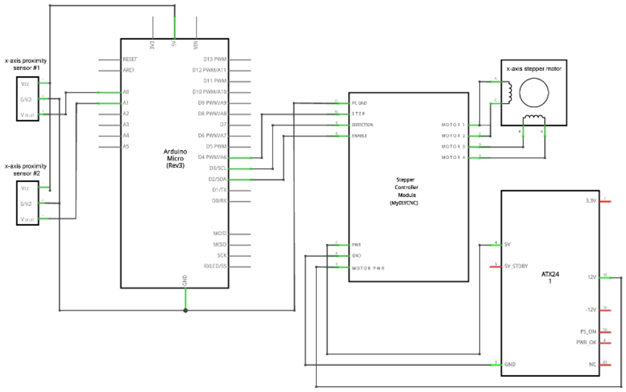

**Circuit diagram of the motor drivers.** This is a representative wiring diagram illustrating the connections for a single axis (the x-axis). The motors and proximity sensors for the other axes can be added via separate stepper controller modules and connected to the open analog input pins (for the sensors) and digital output pins (for the motor controllers).

#### 1.2 Creating the imaging and illumination system

To record the populations, we utilize a low-cost CMOS camera (#1) coupled with a 10X zoom lens (#2) to modify the camera's magnification and field of view in order to observe an entire device. The lens is then mounted on the HDPE mount (#3). To increase video contrast, we machined a collimator (#5) that is tension mounted with screws on the end of the zoom lens. To reduce the amount of light exposure on organisms, we have an automated LED illumination source that automatically switches on when recording behavioral videos or by control of the experimenter via the GUI. The LED source is comprised of two concentric red LED rings (#6-7). The LED rings are mounted on a custom made, laser-cut acrylic LED holder (#8) and is tension mounted via screws on the end of the collimator. See below for an image of the imaging and illumination system. The LEDs are wired to plug connectors (#16-17) and are controlled by a microcontroller connected to computer USB port (#10-11, 15). The system is powered via a 12V power adapter (#12-14) and is contained within a custom machined project box (4.724" L x 3.157" W X 2.311", #9).

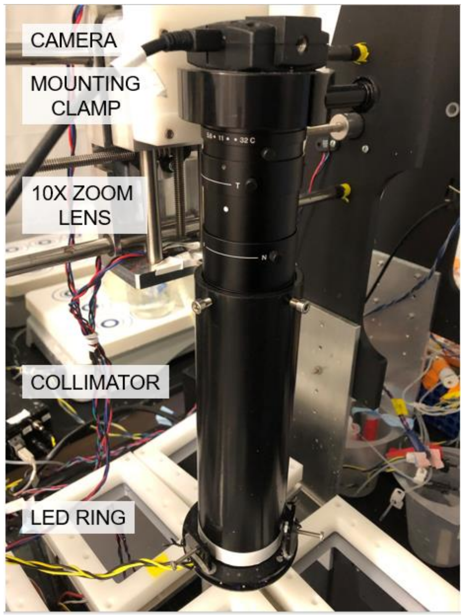

**Close up view of the imaging components.**
Major components are labeled.

### 2. Temperature Control System

Temperature plays a significant role in the development and aging process of *C. elegans*. As a result, it is crucial to provide and maintain precise thermal control over the population. To enable modularity and flexibility for different thermal conditions we developed small thermal modules that can hold up to 4 microfluidic devices. The circuit to control the thermal modules is modular, with a single circuit corresponding to a single thermal module. As a result, it is easy to scale the number of modules according to the experimental need. The BOM assumes the creation of 6 temperature modules, which is the number of modules that can comfortably fit on the platform. However, based on the size of the platform, that can be easily scaled. The following sections detail how to build a single thermal module, along with its corresponding circuitry.

#### 2.1 Building the thermal module

To provide precise heating and cooling capabilities, we used Peltier plates (#24) embedded within machined thermal modules. The module is composed of several layers. A thin copper plate (4.25”L x 3”W x 0.125”, #27) is in thermal contact with a Peltier module, allowing for consistent thermal distribution across a larger surface. To allow for dark-field imaging via reflected light from the over LED ring illumination, a reflective silicon surface (#37) is layered above the copper plate. The Peltier is insulated by a layer of foam (#30) to reduce unintended thermal loss. To prevent overheating and for improved heat dissipation the opposite side of the Peltier is flush with an aluminum plate (5.25”L x 4”W x 0.25”, #29). The plate is attached to a copper heat sink and cooling fan (#2, 25). The Peltier, thermocouple, and cooling fan wiring extend and connect to the main temperature controller via plug connectors (#16, 23). The layers are compression mounted together via screws (#36), which are coupled with machined plastic coverings (#32) and rubber feet (#33) to support and stabilize each module on the platform surface.  A machined plastic outer casing (#28) aids in the compression and allows for long-term mounting of microfluidic devices on the module via screwed clamps (#31, 34-35). A thin layer of thermal paste (#38) is between all layers of the module for improved heat transfer. The CAD files for all machined parts can be found on GrabCAD. See below for images of the thermal module and of microfluidic devices mounted on the module.

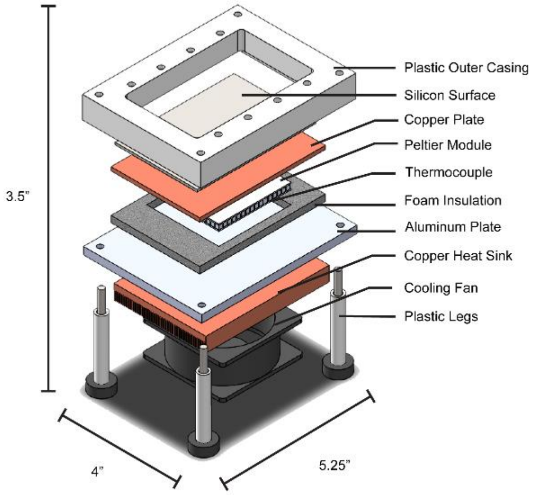

**Exploded view of the thermal module.**

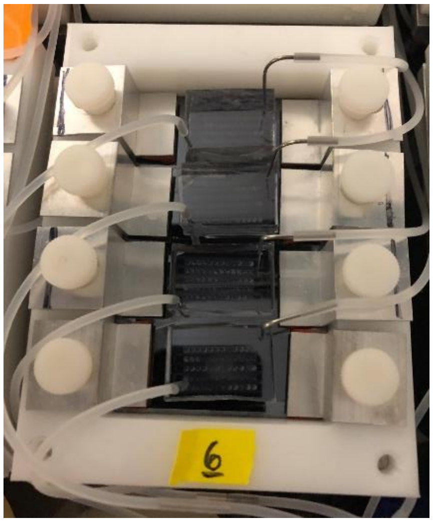

**Microfluidic devices clamped onto the thermal module**. The plastic outer casing provides threads for the aluminum machined clamps to be screwed into, providing pressure to clamp the devices in place over long periods of time.

2.2 Creation the temperature control circuit

The temperature control circuit needs to be able to read analog temperature values from the thermocouple embedded within the temperature module and adjust outputted voltages to account for changes in the set value or ambient temperatures. To do so, we use a basic voltage divider circuit to read resistance changes in the thermistor. The values are reported to the microcontroller, connected to a computer via USB port, and converted via a look-up table into temperature values (#15, 20-21). The required output is then calculated via PID controller, which alters the output magnitude (via PWM analog output) and direction (via digital output) using a DPDT relay, thus allowing heating and/or cooling to reach the desired set point. Due to the high current requirements of the Peltier, we incorporated a voltage buffering state after the PWM output (#3-10). The circuit diagram is shown below and is soldered onto prototyping breadboards (#12). Each circuit was wired to the thermal module via plug connectors (#16-17, 23). The entire system is powered by an external power supply (#18-19,22). All breadboards are contained and fastened (#12) within a custom machined project box (8"L x 12"W X 4 ", #1). For added heat dissipation of the circuits, an array of fans (#2) are embedded within the sides of the enclosure, shown below.

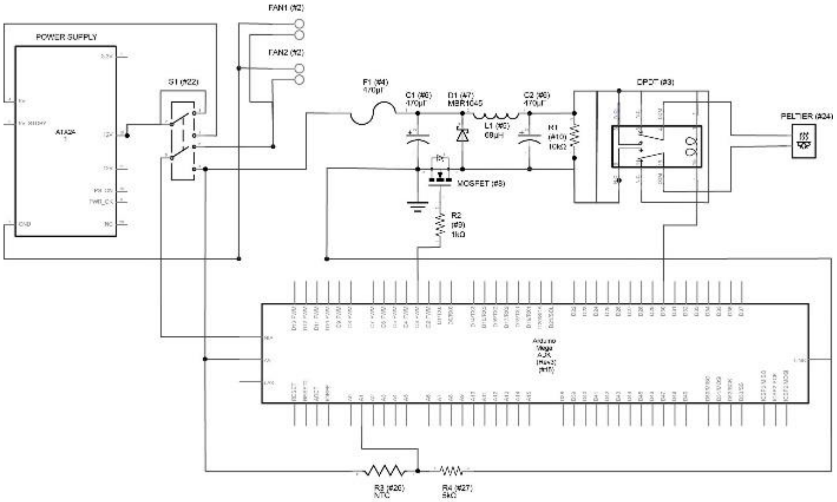

**Circuit diagram for the temperature control circuit.** This is a representative example of a single temperature controller module. It can be easily adapted for more controllers by increasing the number of modular circuits and thermal modules connected to the power supply and microcontroller.

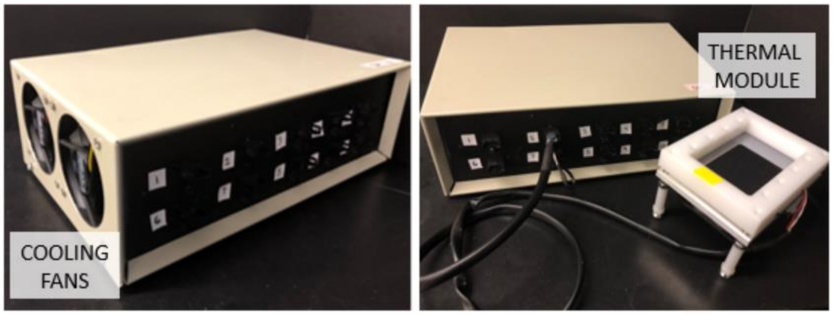

**Temperature Controller System.** (*left*) Temperature control box with cooling side fans. (*right*) Control box with thermal module

**3. Fluid Flow System**

Maintaining a constant food level across the entirety of the worm's lifespan and delivering a wide range of food levels on-chip is largely depending on being able to ensure a constant fluid flow over time to maintain desired food levels within the device. As a result, there is a need to 1) deliver a consistent driving pressure over time for pressure-based fluid flow, and 2) reduce bacterial accumulation and clogging within the system to ensure constant flow. To do so, we have developed two different systems to ensure robust fluid flow and culture over long periods of time.

3.1 Pressure Delivery System

To ensure automated food delivery and we must be able to set and deliver a consistent pressure value over long period of time. Furthermore, to automatically load the microfluidic devices (the process of which is detailed in prior work [25]), it is necessary to quickly apply and release a variety of pressures and activate off-chip solenoid pinch valves to start and stop flow on-chip. To accomplish these needs, we developed a pressure delivery system which acts as an integrated pressure controller system to incorporate independent pressure controls, pneumatic valves, and accessory outlets to drive components such as solenoid pinch valves to control flow off-chip. See below for an image of the system.

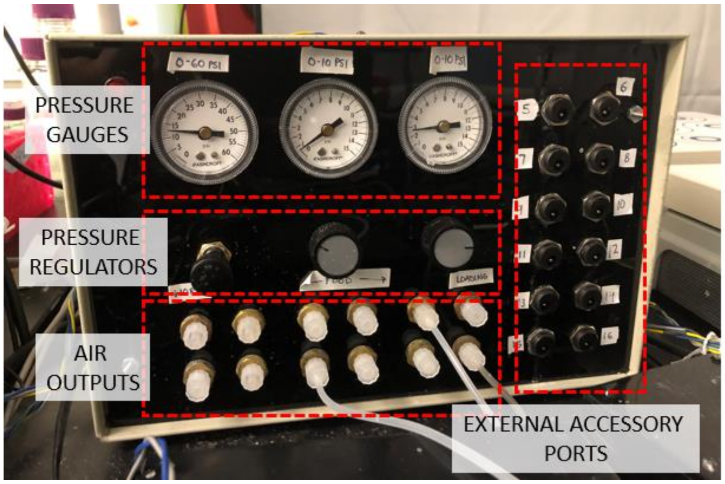

**Pressure delivery system.** User controls (ex. pressure regulators) are highlighted and labeled.

Pressure from an in-house air source is inputted to the system and split across three sets of pressure regulators, with ranges from 0-15PSI to 0-60PSI (#9, 12-14, 16, 18-20). The set pressures are easily modulated and the values are indicated via pressure gauges (#10-11). Two of the regulators are attached to pneumatic valves, allowing electronic switching between the regulated input value and atmospheric value for rapid pressurization/depressurization (#1-3). One directly connects the pressure output to the regulated input for constant pressure-flow applications. Outputted air flow is connected to pressurized reservoirs via tubing with luer connectors (#15, 17, 21). To easily control the valves and accessory power outputs, they are connected to a USB-controllable LED controller, which controls of all electronic outputs to a single USB connection to a computer (#5-8). It is powered via an external power supply (#22, 24, 26), and has panel mounts for external accessory components, such as solenoid pinch valves (#23, 25, 27-29). The parts are housed in a custom-machined project box (11.5”L x 7.5”W x 7”H, #4, 30). Shown below are diagrams of the air flow the electronic circuit diagram.

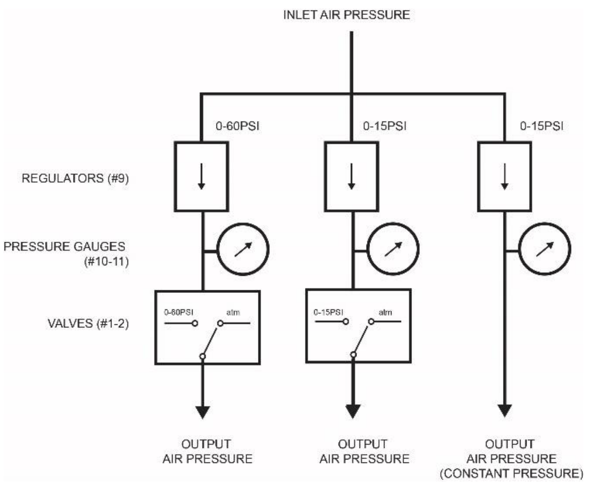

**Pneumatic diagram of the pressure delivery system**

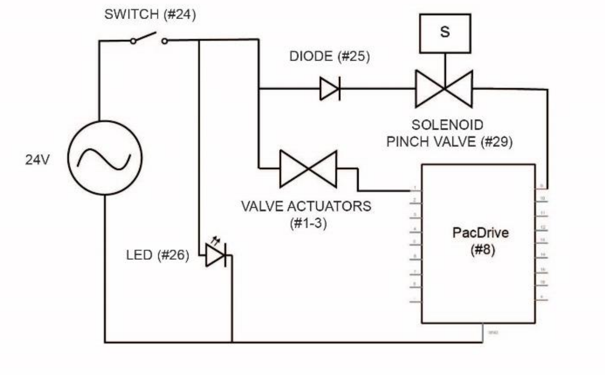

**Circuit diagram for the pressure box.** This is a representative circuit showing the use of one actuating valve and one solenoid pinch valve. More can be easily added in parallel and connected to separate pins in the PacDrive controller.

#### 3.2 Clog Detection System

Even with in-line filters and constant bacterial agitation within the reservoir via stir bar, high food concentrations can create bacterial accumulation in the upstream tubing and filter. This impacts the flow rate within the device and can lead to clogging. To prevent that, we engineered a continuous pressure monitoring device that is able to sense significant pressure drops which signal the presence of clogging upstream. The system then notifies the experimenter via text so that certain parts, such as filters, can be changed before the formation of clogs.

Miniature pressure sensors (#4) are wired as part of an amplified Wheatstone bridge circuit (#5-7). Changes in voltage are read by a microcontroller (#1) and reported in the LabVIEW GUI. The sensors are connected to the system via plug connectors (#2-3) and are powered by an external 12V supply (#8-9). The circuit is soldered to a prototyping breadboard (#10). It is housed in the same project-box as the pressure delivery system. See below for the electronic circuit diagram.

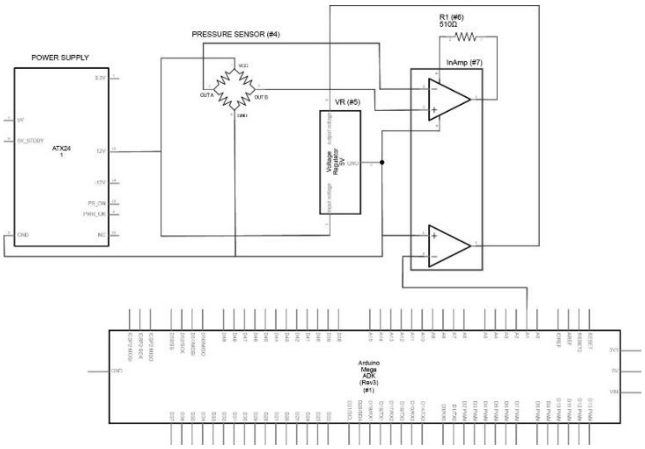

**Circuit diagram of the clog detection system**. This is a representative circuit showing the connections for a single sensor. More sensors can be added and connected to different analog input pins of the microcontroller.

**HeALTH Bill of Materials**

Below is a complete list of materials used to create HeALTH. This list assumes the creation of 6 temperature modules, which is the number of modules that can comfortably fit on the platform, along with 6 pressure sensors and solenoid valves, for each set of devices on a thermal module. Items are organized by subsystem and are numbered accordingly within the section. The prices and items are current as of June 2018; however, availability and pricing could be subject to change. Notes, such as whether the part needs to be machined, are included as well.

| | Item # | Item Name/Description | Vendor | Part Number/ID | Number | Unit Price | Total Cost | Notes |
|---|---|---|---|---|---|---|---|---|
| Imaging System | 1 | DCC1545M - USB 2.0 CMOS Camera | ThorLabs | DCC1545M | 1 | 376.62 | 376.62 | |
| | 2 | 10X (13 - 130mm FL) C-Mount, Close Focus Zoom lens | Edmund Optics | #54-363 | 1 | 595 | 595 | |
| | 3 | 10X Zoom Lens Mounting Clamp | Edmund Optics | #56-547 | 1 | 60 | 60 | |
| | 4 | HDPE mount (connecting lens mounting clamp to the CNC system) | McMaster Carr | 8619K613 | 1 | 11.05 | 11.05 | needs to be machined (see CAD file) |
| | 5 | Acetal Resin Tube collimator | McMaster Carr | 1830T289 | 1 | 15.32 | 15.32 | needs to be machined (see CAD file) |
| | 6 | Red LED Ring (60mm Diameter) | SuperBrightLEDs | AE60-R30-BK | 1 | 7.95 | 7.95 | |
| | 7 | Red LED Halo Ring (80mm Diameter) | SuperBrightLEDs | AE80-R48-BK | 1 | 11.95 | 11.95 | |
| | 8 | Acrylic LED Ring Mount (connecting LED rings to collimator) | McMaster Carr | 8505K741 | 1 | 7.14 | 7.14 | needs to be machined (see CAD file) |
| | 9 | Project Box for LED Control (4.724" L x 3.157" W (119.99mm x 80.19mm) X 2.311" (58.70mm)) | Digikey | HM110-ND | 1 | 6.5 | 6.5 | needs to be machined |
| | 10 | PacDrive USB driver board Type: Special (Outputs OFF at power on) | Ultimarc | PPACDSP+LU04 | 1 | 30 | 30 | |
| | 11 | USB 2.0 Cable (A Male to B Male) | Digikey | 1175-1088-ND | 2 | 3.25 | 6.5 | |
| | 12 | Toggle Switch (SPDT) | Digikey | EG2350-ND | 1 | 2.43 | 2.43 | |
| | 13 | 12V AC/DC Wall Power Adapter | Digikey | 62-1230-ND | 1 | 12.21 | 12.21 | |

| | | | | | | | | |
|---|---|---|---|---|---|---|---|---|
| | 14 | Power Barrel Connector Jack | Digikey | EJ508A-ND | 1 | 1.38 | 1.38 | |
| | 15 | USB Type B - Type A Adapter | Allied Electronics | 70126196 | 2 | 10.05 | 20.1 | |
| | 16 | 4 wire plug connector pin (male) | Digikey | 3M155973-ND | 7 | 3.22 | 22.54 | |
| | 17 | 4 wire plug connector pin (female) | Digikey | 3M156151-ND | 7 | 3.5 | 24.5 | |
| | 18 | Aluminum platform sides | McMaster Carr | 8975K432 | 1 | 23.18 | 23.18 | needs to be machined (see CAD file) |
| | 19 | Acrylic Platform | McMaster Carr | 8505K731 | 1 | 33.13 | 33.13 | needs to be machined (see CAD file) |
| | 20 | Hall Effect Magnetic Sensors | Mouser | 934-551003H02A | 6 | 6.88 | 41.28 | |
| | 21 | DIY CNC kit | MyDIYCNC | MyDIYCNC Desktop CNC Machine | 1 | 445 | 445 | Exact system is currently discontinued, but should be able to adapt to other, common DIY CNC kits |
| | 22 | Project Box for Motor Control ( 6.112" L x 4.612" W (155.24mm x 117.14mm) X 2.356" (59.84mm)) | Digikey | 377-1218-ND | 1 | 9.7 | 9.7 | needs to be machined |
| | 23 | Arduino Micro | Arduino | A000053 | 1 | 19.8 | 19.8 | |
| | 24 | 3 wire plug connector pin (male) | Digikey | 3M155968-ND | 1 | 2.72 | 2.72 | |
| | 25 | 3 wire plug connector pin (female) | Digikey | 3M155845-ND | 1 | 2.92 | 2.92 | |
| | 26 | 4 position terminal block for attaching wire to board | Digikey | A98335-ND | 4 | 2.68 | 10.72 | |
| | 27 | 2 position terminal block for attaching wire to board | Digikey | A98333-ND | 2 | 0.88 | 1.76 | |
| | 28 | Solder prototyping breadboard | Digikey | SBB400-ND | 1 | 3.99 | 3.99 | |
| **Temperature Control System** | 1 | Project Box for Temperature Control (8.000" L x 12.000" W (203.20mm x 304.80mm) X 4.000" (101.60mm)) | Digikey | HM302-ND | 1 | 60.5 | 60.5 | needs to be machined |
| | 2 | CPU Cooling Fan | Amazon | R4-S8R-20AK-GP | 10 | 5.71 | 57.1 | |

| | | | | | | | | |
|---|---|---|---|---|---|---|---|---|
| | 3 | DPDT relay | Digikey | PB383-ND | 6 | 2.36 | 14.16 | |
| | 4 | On-board 4A fuse | Digikey | F2344CT-ND | 6 | 1.842 | 11.052 | |
| | 5 | 68 uH inductor | Digikey | AIAP-03-680K-ND | 6 | 1.55 | 9.3 | |
| | 6 | 470 uF capacitor | Digikey | P16299-ND | 12 | 0.941 | 11.292 | |
| | 7 | Schottky diode | Digikey | MBR1045PBF-ND | 6 | 1.1 | 6.6 | |
| | 8 | logic level MOSFET | Digikey | 2SK4017Q-ND | 6 | 0.67 | 4.02 | |
| | 9 | 1 kohm resistor | Digikey | PPC1.0KBCT-ND | 6 | 0.44 | 2.64 | |
| | 10 | 10 kohm resistor | Digikey | PPC10KBCT-ND | 6 | 0.44 | 2.64 | |
| | 11 | Solder Prototyping Breadboard | Digikey | SBB206-ND | 6 | 2.49 | 14.94 | |
| | 12 | Breadboard Adhesive Fastener | Digikey | RPC1096-ND | 12 | 0.932 | 11.184 | |
| | 13 | 2 position terminal block for attaching wire to board | Digikey | A98333-ND | 6 | 0.88 | 5.28 | |
| | 14 | 4 position terminal block for attaching wire to board | Digikey | A98335-ND | 6 | 2.68 | 16.08 | |
| | 15 | Arduino MEGA | Arduino | A000067 | 1 | 38.5 | 38.5 | |
| | 16 | 4 wire plug connector pin (male) | Digikey | 3M155973-ND | 6 | 3.22 | 19.32 | |
| | 17 | 4 wire plug connector pin (female) | Digikey | 3M156151-ND | 6 | 3.5 | 21 | |
| | 18 | Power Barrel Connector | Digikey | CP-011B-ND | 2 | 1.6 | 3.2 | |
| | 19 | Cable assembly (to aux power) | Digikey | CP-2188-ND | 2 | 2.5 | 5 | |
| | 20 | USB Type B - Type A Adapter | Allied Electronics | 70126196 | 1 | 10.05 | 10.05 | |
| | 21 | USB 2.0 Cable (A Male to B Male) | Digikey | 1175-1088-ND | 1 | 3.25 | 3.25 | |
| | 22 | Toggle Switch (DPDT) | Digikey | EG2398-ND | 1 | 3.25 | 3.25 | |
| | 23 | 2-pin JST SM Plug + Receptacle Cable Set | Adafruit | 2880 | 6 | 0.75 | 4.5 | |
| | 24 | Peltier Plate | Digikey | 926-1267-ND | 6 | 65.92 | 395.52 | |
| | 25 | CPU Heat Sink | Newegg | N82E16835114039 | 6 | 25.75 | 154.5 | |
| | 26 | Surface Thermistor (10kohm) | Omega | SA2F-TH-44031-40 | 6 | 68.5 | 411 | |
| | 27 | Precision Resistor (5kohm) | Digikey | 696-1566-ND | 6 | 6.12 | 36.72 | |
| | 28 | Copper Plate | McMaster | 8963K703 | 1 | 45.19 | 45.19 | needs to be machined (see CAD file) |

| | | | | | | | | |
|---|---|---|---|---|---|---|---|---|
| | 29 | HDPE Plastic Casing | McMaster | 8619K482 | 1 | 34.31 | 34.31 | needs to be machined (see CAD file) |
| | 30 | Aluminum Plate | McMaster | 8975K437 | 1 | 34.8 | 34.8 | needs to be machined (see CAD file) |
| | 31 | Foam Insulation | McMaster | 5541K11 | 1 | 10.62 | 10.62 | |
| | 32 | Plastic screws | McMaster | 94323A837 | 1 | 9.93 | 9.93 | |
| | 33 | Plastic legs | McMaster | 8624K12 | 1 | 6.72 | 6.72 | needs to be machined (see CAD file) |
| | 34 | Rubber feet | McMaster | 9541K2 | 1 | 12.48 | 12.48 | |
| | 35 | Aluminum bracket | McMaster | 7062T13 | 1 | 7.9 | 7.9 | needs to be machined (see CAD file) |
| | 36 | Bracket rubber seal | McMaster | 1129A989 | 1 | 8.75 | 8.75 | |
| | 37 | 18-8 Screws | McMaster | 92196A540 | 1 | 9.53 | 9.53 | |
| | 38 | Silicon Wafer | University Wafer | 857 | 6 | 23.9 | 143.4 | |
| | 39 | Thermal Paste | Amazon | AS5-3.5G | 1 | 6 | 6 | |
| **Pressure Delivery System** | 1 | Valve (set of 4) actuator | KSCDirect | ASC 18800056 | 0.5 | 278.75 | 139.375 | |
| | 2 | Valve (set of 2) actuator | KSCDirect | ASC 18800054 | 0.5 | 145.02 | 72.51 | |
| | 3 | Valve Connector | KSCDirect | ASC 88118802 | 1 | 11.3 | 11.3 | |
| | 4 | Project Box (for Pressure Box) (11.5"x7.5"x7") | Allied Electronics | 70164768 | 1 | 103.45 | 103.45 | needs to be machined |
| | 5 | USB Type B - Type A Adapter | Allied Electronics | 70126196 | 2 | 10.05 | 20.1 | |
| | 6 | USB 2.0 Cable (A Male to B Male) | Digikey | 1175-1088-ND | 2 | 3.25 | 6.5 | |
| | 7 | USB 2.0 Cable (A Male to B Male) | Digikey | 1597-1533-ND | 2 | 1.99 | 3.98 | |
| | 8 | PacDrive USB driver board Type: Special (Outputs OFF at power on) | Ultimarc | PPACDSP+LU04 | 1 | 30 | 30 | |
| | 9 | Regulator (0.5-60 PSI) for valves | McMaster | 43275K14 | 3 | 40 | 120 | |
| | 10 | Pressure gauge (0-60 PSI) for valves | McMaster | 3846K43 | 1 | 12.39 | 12.39 | |
| | 11 | Pressure gauge (0-15 PSI) for valves | McMaster | 3846K41 | 2 | 10.46 | 20.92 | |
| | 12 | Polybutylene & Brass Push-to-Connect Fitting Adapter from pressure gauges to push to connect fitting | McMaster | 5111K664 | 3 | 3.57 | 10.71 | |

| | | | | | | | |
|---|---|---|---|---|---|---|---|
| | 13 | Polybutylene & Nickel-Pltd Brass adapter from pressure regulators and valve manifold to push to connect fitting | McMaster | 52065K124 | 6 | 2.35 | 14.1 |
| | 14 | Y airway splitter | McMaster | 5111K402 | 2 | 5.1 | 10.2 |
| | 15 | Miniature Brass Coupler to external outlets | McMaster | 5454K85 | 1 | 10.31 | 10.31 |
| | 16 | Polybutylene & Brass Push-to-Connect Fitting Tee airway splitter | McMaster | 5111K172 | 3 | 3.4 | 10.2 |
| | 17 | Nylon and Nickel-Plated Brass Tube Fitting Adapter for couples valve manifold to external outlets | McMaster | 5779K244 | 6 | 2.76 | 16.56 |
| | 18 | Polybutylene & Brass panel mount coupler to external pressure source | McMaster | 5111K202 | 1 | 9.3 | 9.3 |
| | 19 | Polybutylene & Brass adapter to external pressure source | McMaster | 5111K92 | 1 | 1.89 | 1.89 |
| | 20 | Nylon tubing for internal air connections | McMaster | 5112K52 | 2 | 0.31 | 0.62 |
| | 21 | Male Luer Integral Lock Ring to 10-32 Special Tapered Thread, Natural Polypropylene | Nordson Medical | XMTLL-6 | 12 | 0.3498 | 4.1976 |
| | 22 | Power supply | DigiKey | 271-2592-ND | 1 | 23.41 | 23.41 |
| | 23 | Panel mount plugs | DigiKey | CP-011B-ND | 10 | 1.44 | 14.4 |
| | 24 | Power switch | DigiKey | 679-1211-ND | 1 | 7.22 | 7.22 |
| | 25 | Protection diode 1A | Digikey | 1N4004RLGO SCT-ND | 10 | 1.87 | 18.7 |
| | 26 | Panel mount LED | Digikey | 350-2101-ND | 1 | 5.24 | 5.24 |
| | 27 | Screw terminal, connects wires | DigiKey | WM15900-ND | 5 | 1.79 | 8.95 |
| | 28 | Cable assumbly (to aux power) | Digikey | CP-2188-ND | 2 | 2.5 | 5 |
| | 29 | Solenoid pinch valve (2-way normally closed) | Cole Parmer | EW-98302-10 | 6 | 99.5 | 597 |
| | 30 | 4-40 Machine Screws | Allied Electronics | 70126097 | 0.1 | 8.92 | 0.892 |

| | | | | | | | |
|---|---|---|---|---|---|---|---|
| Clog Detection System | 1 | Arduino MEGA | Arduino | A000067 | 1 | 38.5 | 38.5 |
| | 2 | 4 wire plug connector pin (male) | Digikey | 3M155973-ND | 6 | 3.22 | 19.32 |
| | 3 | 4 wire plug connector pin (female) | Digikey | 3M156151-ND | 6 | 3.5 | 21 |
| | 4 | Pressure sensors (15 PSI) | Mouser | 785-24PCCFG6G | 6 | 41.65 | 249.9 |
| | 5 | 5V voltage regulator | Mouser | 511-L7805CV | 1 | 10.05 | 10.05 |
| | 6 | 510ohm resistor | Mouser | 667-ERG-1SJ511 | 6 | 3.25 | 19.5 |
| | 7 | Instrumentation amplifier | Mouser | 595-INA126PA | 6 | 3.15 | 18.9 |
| | 8 | 3 wire plug connector pin (male) | Digikey | 3M155968-ND | 1 | 2.72 | 2.72 |
| | 9 | 3 wire plug connector pin (female) | Digikey | 3M155845-ND | 1 | 2.92 | 2.92 |
| | 10 | Solder Prototyping Breadboard | Digikey | SBB206-ND | 1 | 2.49 | 2.49 |
| Misc. Items | 1 | Multi-position magnetic stirrers | VWR | 12621-046 | 1 | 1609 | 1609 |
| | 2 | Power supply | Newegg | N82E16817438018 | 1 | 119.99 | 119.99 |
| | 3 | Cable set | Newegg | 9SIAA6P4G38262 | 1 | 49.9 | 49.9 |
| | 4 | Heat shrink tubing | Amazon | B075WR9FVL | 1 | 7.99 | 7.99 |
| | 5 | Wire | Digikey | C2016B-100-ND | 1 | 18.7 | 18.7 |

**Supplementary Note 2.** LASSO regression

Below is a list of behavioral metrics that were extracted from the data set. This is by no means an exhaustive list of potential extracted behavioral metrics, but simply what we used to perform LASSO regression to observe how stereotyped lifespan is across a population in relation to behavioral decline over time. Bolded variables indicated variables actually inputted into the LASSO regression model for the wild-type population cultured at 25°C with food levels of $OD_{600}$5. Other variables were censored due to data sparsity. Additionally, Day 0 indicates activity at the L4 larval stage.

**'ampData_maxVal_maxValDay_Day0'**
**'ampData_maxVal_maxValDay_Day1'**
**'ampData_maxVal_maxValDay_Day2'**
**'ampData_maxVal_maxValDay_Day3'**
**'ampData_maxVal_maxValDay_Day4'**
**'ampData_maxVal_maxValDay_Day5'**
**'ampData_maxVal_maxValDay_Day6'**
**'ampData_maxVal_maxValDay_Day7'**
**'ampData_maxVal_maxValDay_Day8'**
**'ampData_maxVal_maxValDay_Day9'**
**'ampData_maxVal_maxValDay_Day10'**
**'ampData_maxVal_avgValDay_Day0'**
**'ampData_maxVal_avgValDay_Day1'**
**'ampData_maxVal_avgValDay_Day2'**
**'ampData_maxVal_avgValDay_Day3'**
**'ampData_maxVal_avgValDay_Day4'**
**'ampData_maxVal_avgValDay_Day5'**
**'ampData_maxVal_avgValDay_Day6'**
**'ampData_maxVal_avgValDay_Day7'**
**'ampData_maxVal_avgValDay_Day8'**
**'ampData_maxVal_avgValDay_Day9'**
**'ampData_maxVal_avgValDay_Day10'**
'ampData_maxVal_maxValDay_high_Day0'
**'ampData_maxVal_maxValDay_high_Day1'**
**'ampData_maxVal_maxValDay_high_Day2'**
**'ampData_maxVal_maxValDay_high_Day3'**
**'ampData_maxVal_maxValDay_high_Day4'**
**'ampData_maxVal_maxValDay_high_Day5'**
**'ampData_maxVal_maxValDay_high_Day6'**
**'ampData_maxVal_maxValDay_high_Day7'**
**'ampData_maxVal_maxValDay_high_Day8'**
**'ampData_maxVal_maxValDay_high_Day9'**
**'ampData_maxVal_maxValDay_high_Day10'**
'ampData_maxVal_avgValDay_high_Day0'
**'ampData_maxVal_avgValDay_high_Day1'**

**'csData_maxVal_maxValDay_Day0'**
**'csData_maxVal_maxValDay_Day1'**
**'csData_maxVal_maxValDay_Day2'**
**'csData_maxVal_maxValDay_Day3'**
**'csData_maxVal_maxValDay_Day4'**
**'csData_maxVal_maxValDay_Day5'**
**'csData_maxVal_maxValDay_Day6'**
**'csData_maxVal_maxValDay_Day7'**
**'csData_maxVal_maxValDay_Day8'**
**'csData_maxVal_maxValDay_Day9'**
**'csData_maxVal_maxValDay_Day10'**
**'csData_maxVal_avgValDay_Day0'**
**'csData_maxVal_avgValDay_Day1'**
**'csData_maxVal_avgValDay_Day2'**
**'csData_maxVal_avgValDay_Day3'**
**'csData_maxVal_avgValDay_Day4'**
**'csData_maxVal_avgValDay_Day5'**
**'csData_maxVal_avgValDay_Day6'**
**'csData_maxVal_avgValDay_Day7'**
**'csData_maxVal_avgValDay_Day8'**
**'csData_maxVal_avgValDay_Day9'**
**'csData_maxVal_avgValDay_Day10'**
**'csData_maxVal_maxValDay_high_Day0'**
**'csData_maxVal_maxValDay_high_Day1'**
**'csData_maxVal_maxValDay_high_Day2'**
**'csData_maxVal_maxValDay_high_Day3'**
**'csData_maxVal_maxValDay_high_Day4'**
**'csData_maxVal_maxValDay_high_Day5'**
'csData_maxVal_maxValDay_high_Day6'
'csData_maxVal_maxValDay_high_Day7'
'csData_maxVal_maxValDay_high_Day8'
'csData_maxVal_maxValDay_high_Day9'
'csData_maxVal_maxValDay_high_Day10'
**'csData_maxVal_avgValDay_high_Day0'**
**'csData_maxVal_avgValDay_high_Day1'**

**'ampData_maxVal_avgValDay_high_Day2'**
**'ampData_maxVal_avgValDay_high_Day3'**
**'ampData_maxVal_avgValDay_high_Day4'**
**'ampData_maxVal_avgValDay_high_Day5'**
**'ampData_maxVal_avgValDay_high_Day6'**
**'ampData_maxVal_avgValDay_high_Day7'**
**'ampData_maxVal_avgValDay_high_Day8'**
**'ampData_maxVal_avgValDay_high_Day9'**
**'ampData_maxVal_avgValDay_high_Day10'**
**'ampData_maxVal_declinePT_duration'**
**'ampData_maxVal_relativeHighPeriod_duration'**
**'ampData_meanVal_maxValDay_Day0'**
**'ampData_meanVal_maxValDay_Day1'**
**'ampData_meanVal_maxValDay_Day2'**
**'ampData_meanVal_maxValDay_Day3'**
**'ampData_meanVal_maxValDay_Day4'**
**'ampData_meanVal_maxValDay_Day5'**
**'ampData_meanVal_maxValDay_Day6'**
**'ampData_meanVal_maxValDay_Day7'**
**'ampData_meanVal_maxValDay_Day8'**
**'ampData_meanVal_maxValDay_Day9'**
**'ampData_meanVal_maxValDay_Day10'**
**'ampData_meanVal_avgValDay_Day0'**
**'ampData_meanVal_avgValDay_Day1'**
**'ampData_meanVal_avgValDay_Day2'**
**'ampData_meanVal_avgValDay_Day3'**
**'ampData_meanVal_avgValDay_Day4'**
**'ampData_meanVal_avgValDay_Day5'**
**'ampData_meanVal_avgValDay_Day6'**
**'ampData_meanVal_avgValDay_Day7'**
**'ampData_meanVal_avgValDay_Day8'**
**'ampData_meanVal_avgValDay_Day9'**
**'ampData_meanVal_avgValDay_Day10'**
'ampData_meanVal_maxValDay_high_Day0'
**'ampData_meanVal_maxValDay_high_Day1'**
**'ampData_meanVal_maxValDay_high_Day2'**
**'ampData_meanVal_maxValDay_high_Day3'**
**'ampData_meanVal_maxValDay_high_Day4'**
**'ampData_meanVal_maxValDay_high_Day5'**
**'ampData_meanVal_maxValDay_high_Day6'**
**'ampData_meanVal_maxValDay_high_Day7'**
**'ampData_meanVal_maxValDay_high_Day8'**
**'ampData_meanVal_maxValDay_high_Day9'**

**'csData_maxVal_avgValDay_high_Day2'**
**'csData_maxVal_avgValDay_high_Day3'**
**'csData_maxVal_avgValDay_high_Day4'**
**'csData_maxVal_avgValDay_high_Day5'**
'csData_maxVal_avgValDay_high_Day6'
'csData_maxVal_avgValDay_high_Day7'
'csData_maxVal_avgValDay_high_Day8'
'csData_maxVal_avgValDay_high_Day9'
'csData_maxVal_avgValDay_high_Day10'
**'csData_maxVal_declinePT_duration'**
**'csData_maxVal_relativeHighPeriod_duration'**
**'csData_meanVal_maxValDay_Day0'**
**'csData_meanVal_maxValDay_Day1'**
**'csData_meanVal_maxValDay_Day2'**
**'csData_meanVal_maxValDay_Day3'**
**'csData_meanVal_maxValDay_Day4'**
**'csData_meanVal_maxValDay_Day5'**
**'csData_meanVal_maxValDay_Day6'**
**'csData_meanVal_maxValDay_Day7'**
**'csData_meanVal_maxValDay_Day8'**
**'csData_meanVal_maxValDay_Day9'**
**'csData_meanVal_maxValDay_Day10'**
**'csData_meanVal_avgValDay_Day0'**
**'csData_meanVal_avgValDay_Day1'**
**'csData_meanVal_avgValDay_Day2'**
**'csData_meanVal_avgValDay_Day3'**
**'csData_meanVal_avgValDay_Day4'**
**'csData_meanVal_avgValDay_Day5'**
**'csData_meanVal_avgValDay_Day6'**
**'csData_meanVal_avgValDay_Day7'**
**'csData_meanVal_avgValDay_Day8'**
**'csData_meanVal_avgValDay_Day9'**
**'csData_meanVal_avgValDay_Day10'**
**'csData_meanVal_maxValDay_high_Day0'**
**'csData_meanVal_maxValDay_high_Day1'**
**'csData_meanVal_maxValDay_high_Day2'**
**'csData_meanVal_maxValDay_high_Day3'**
'csData_meanVal_maxValDay_high_Day4'
'csData_meanVal_maxValDay_high_Day5'
'csData_meanVal_maxValDay_high_Day6'
'csData_meanVal_maxValDay_high_Day7'
'csData_meanVal_maxValDay_high_Day8'
'csData_meanVal_maxValDay_high_Day9'

**'ampData_meanVal_maxValDay_high_Day10'**
'ampData_meanVal_avgValDay_high_Day0'
**'ampData_meanVal_avgValDay_high_Day1'**
**'ampData_meanVal_avgValDay_high_Day2'**
**'ampData_meanVal_avgValDay_high_Day3'**
**'ampData_meanVal_avgValDay_high_Day4'**
**'ampData_meanVal_avgValDay_high_Day5'**
**'ampData_meanVal_avgValDay_high_Day6'**
**'ampData_meanVal_avgValDay_high_Day7'**
**'ampData_meanVal_avgValDay_high_Day8'**
**'ampData_meanVal_avgValDay_high_Day9'**
**'ampData_meanVal_avgValDay_high_Day10'**
**'ampData_meanVal_declinePT_duration'**
**'ampData_meanVal_relativeHighPeriod_duration'**
**'dpData_maxVal_maxValDay_Day0'**
**'dpData_maxVal_maxValDay_Day1'**
**'dpData_maxVal_maxValDay_Day2'**
**'dpData_maxVal_maxValDay_Day3'**
**'dpData_maxVal_maxValDay_Day4'**
**'dpData_maxVal_maxValDay_Day5'**
**'dpData_maxVal_maxValDay_Day6'**
**'dpData_maxVal_maxValDay_Day7'**
**'dpData_maxVal_maxValDay_Day8'**
**'dpData_maxVal_maxValDay_Day9'**
**'dpData_maxVal_maxValDay_Day10'**
**'dpData_maxVal_avgValDay_Day0'**
**'dpData_maxVal_avgValDay_Day1'**
**'dpData_maxVal_avgValDay_Day2'**
**'dpData_maxVal_avgValDay_Day3'**
**'dpData_maxVal_avgValDay_Day4'**
**'dpData_maxVal_avgValDay_Day5'**
**'dpData_maxVal_avgValDay_Day6'**
**'dpData_maxVal_avgValDay_Day7'**
**'dpData_maxVal_avgValDay_Day8'**
**'dpData_maxVal_avgValDay_Day9'**
**'dpData_maxVal_avgValDay_Day10'**
**'dpData_maxVal_maxValDay_high_Day0'**
**'dpData_maxVal_maxValDay_high_Day1'**
**'dpData_maxVal_maxValDay_high_Day2'**
**'dpData_maxVal_maxValDay_high_Day3'**
**'dpData_maxVal_maxValDay_high_Day4'**
'dpData_maxVal_maxValDay_high_Day5'
'dpData_maxVal_maxValDay_high_Day6'

'csData_meanVal_maxValDay_high_Day10'
**'csData_meanVal_avgValDay_high_Day0'**
**'csData_meanVal_avgValDay_high_Day1'**
**'csData_meanVal_avgValDay_high_Day2'**
**'csData_meanVal_avgValDay_high_Day3'**
'csData_meanVal_avgValDay_high_Day4'
'csData_meanVal_avgValDay_high_Day5'
'csData_meanVal_avgValDay_high_Day6'
'csData_meanVal_avgValDay_high_Day7'
'csData_meanVal_avgValDay_high_Day8'
'csData_meanVal_avgValDay_high_Day9'
'csData_meanVal_avgValDay_high_Day10'
**'csData_meanVal_declinePT_duration'**
**'csData_meanVal_relativeHighPeriod_duration'**
**'freqData_meanVal_maxValDay_Day0'**
**'freqData_meanVal_maxValDay_Day1'**
**'freqData_meanVal_maxValDay_Day2'**
**'freqData_meanVal_maxValDay_Day3'**
**'freqData_meanVal_maxValDay_Day4'**
**'freqData_meanVal_maxValDay_Day5'**
**'freqData_meanVal_maxValDay_Day6'**
**'freqData_meanVal_maxValDay_Day7'**
**'freqData_meanVal_maxValDay_Day8'**
**'freqData_meanVal_maxValDay_Day9'**
**'freqData_meanVal_maxValDay_Day10'**
**'freqData_meanVal_avgValDay_Day0'**
**'freqData_meanVal_avgValDay_Day1'**
**'freqData_meanVal_avgValDay_Day2'**
**'freqData_meanVal_avgValDay_Day3'**
**'freqData_meanVal_avgValDay_Day4'**
**'freqData_meanVal_avgValDay_Day5'**
**'freqData_meanVal_avgValDay_Day6'**
**'freqData_meanVal_avgValDay_Day7'**
**'freqData_meanVal_avgValDay_Day8'**
**'freqData_meanVal_avgValDay_Day9'**
**'freqData_meanVal_avgValDay_Day10'**
**'freqData_meanVal_maxValDay_high_Day0'**
**'freqData_meanVal_maxValDay_high_Day1'**
**'freqData_meanVal_maxValDay_high_Day2'**
**'freqData_meanVal_maxValDay_high_Day3'**
**'freqData_meanVal_maxValDay_high_Day4'**
**'freqData_meanVal_maxValDay_high_Day5'**
**'freqData_meanVal_maxValDay_high_Day6'**

'dpData_maxVal_maxValDay_high_Day7'
'dpData_maxVal_maxValDay_high_Day8'
'dpData_maxVal_maxValDay_high_Day9'
'dpData_maxVal_maxValDay_high_Day10'
**'dpData_maxVal_avgValDay_high_Day0'**
**'dpData_maxVal_avgValDay_high_Day1'**
**'dpData_maxVal_avgValDay_high_Day2'**
**'dpData_maxVal_avgValDay_high_Day3'**
**'dpData_maxVal_avgValDay_high_Day4'**
'dpData_maxVal_avgValDay_high_Day5'
'dpData_maxVal_avgValDay_high_Day6'
'dpData_maxVal_avgValDay_high_Day7'
'dpData_maxVal_avgValDay_high_Day8'
'dpData_maxVal_avgValDay_high_Day9'
'dpData_maxVal_avgValDay_high_Day10'
**'dpData_maxVal_declinePT_duration'**
**'dpData_maxVal_relativeHighPeriod_duration'**
**'dpData_meanVal_maxValDay_Day0'**
**'dpData_meanVal_maxValDay_Day1'**
**'dpData_meanVal_maxValDay_Day2'**
**'dpData_meanVal_maxValDay_Day3'**
**'dpData_meanVal_maxValDay_Day4'**
**'dpData_meanVal_maxValDay_Day5'**
**'dpData_meanVal_maxValDay_Day6'**
**'dpData_meanVal_maxValDay_Day7'**
**'dpData_meanVal_maxValDay_Day8'**
**'dpData_meanVal_maxValDay_Day9'**
**'dpData_meanVal_maxValDay_Day10'**
**'dpData_meanVal_avgValDay_Day0'**
**'dpData_meanVal_avgValDay_Day1'**
**'dpData_meanVal_avgValDay_Day2'**
**'dpData_meanVal_avgValDay_Day3'**
**'dpData_meanVal_avgValDay_Day4'**
**'dpData_meanVal_avgValDay_Day5'**
**'dpData_meanVal_avgValDay_Day6'**
**'dpData_meanVal_avgValDay_Day7'**
**'dpData_meanVal_avgValDay_Day8'**
**'dpData_meanVal_avgValDay_Day9'**
**'dpData_meanVal_avgValDay_Day10'**
**'dpData_meanVal_maxValDay_high_Day0'**
**'dpData_meanVal_maxValDay_high_Day1'**
**'dpData_meanVal_maxValDay_high_Day2'**
**'dpData_meanVal_maxValDay_high_Day3'**

**'freqData_meanVal_maxValDay_high_Day7'**
**'freqData_meanVal_maxValDay_high_Day8'**
**'freqData_meanVal_maxValDay_high_Day9'**
**'freqData_meanVal_maxValDay_high_Day10'**
**'freqData_meanVal_avgValDay_high_Day0'**
**'freqData_meanVal_avgValDay_high_Day1'**
**'freqData_meanVal_avgValDay_high_Day2'**
**'freqData_meanVal_avgValDay_high_Day3'**
**'freqData_meanVal_avgValDay_high_Day4'**
**'freqData_meanVal_avgValDay_high_Day5'**
**'freqData_meanVal_avgValDay_high_Day6'**
**'freqData_meanVal_avgValDay_high_Day7'**
**'freqData_meanVal_avgValDay_high_Day8'**
**'freqData_meanVal_avgValDay_high_Day9'**
**'freqData_meanVal_avgValDay_high_Day10'**
**'freqData_meanVal_declinePT_duration'**
**'freqData_meanVal_relativeHighPeriod_duration'**
**'freqData_maxVal_maxValDay_Day0'**
**'freqData_maxVal_maxValDay_Day1'**
**'freqData_maxVal_maxValDay_Day2'**
**'freqData_maxVal_maxValDay_Day3'**
**'freqData_maxVal_maxValDay_Day4'**
**'freqData_maxVal_maxValDay_Day5'**
**'freqData_maxVal_maxValDay_Day6'**
**'freqData_maxVal_maxValDay_Day7'**
**'freqData_maxVal_maxValDay_Day8'**
**'freqData_maxVal_maxValDay_Day9'**
**'freqData_maxVal_maxValDay_Day10'**
**'freqData_maxVal_avgValDay_Day0'**
**'freqData_maxVal_avgValDay_Day1'**
**'freqData_maxVal_avgValDay_Day2'**
**'freqData_maxVal_avgValDay_Day3'**
**'freqData_maxVal_avgValDay_Day4'**
**'freqData_maxVal_avgValDay_Day5'**
**'freqData_maxVal_avgValDay_Day6'**
**'freqData_maxVal_avgValDay_Day7'**
**'freqData_maxVal_avgValDay_Day8'**
**'freqData_maxVal_avgValDay_Day9'**
**'freqData_maxVal_avgValDay_Day10'**
**'freqData_maxVal_maxValDay_high_Day0'**
**'freqData_maxVal_maxValDay_high_Day1'**
**'freqData_maxVal_maxValDay_high_Day2'**
**'freqData_maxVal_maxValDay_high_Day3'**

'dpData_meanVal_maxValDay_high_Day4'
'dpData_meanVal_maxValDay_high_Day5'
'dpData_meanVal_maxValDay_high_Day6'
'dpData_meanVal_maxValDay_high_Day7'
'dpData_meanVal_maxValDay_high_Day8'
'dpData_meanVal_maxValDay_high_Day9'
'dpData_meanVal_maxValDay_high_Day10'
**'dpData_meanVal_avgValDay_high_Day0'**
**'dpData_meanVal_avgValDay_high_Day1'**
**'dpData_meanVal_avgValDay_high_Day2'**
**'dpData_meanVal_avgValDay_high_Day3'**
'dpData_meanVal_avgValDay_high_Day4'
'dpData_meanVal_avgValDay_high_Day5'
'dpData_meanVal_avgValDay_high_Day6'
'dpData_meanVal_avgValDay_high_Day7'
'dpData_meanVal_avgValDay_high_Day8'
'dpData_meanVal_avgValDay_high_Day9'
'dpData_meanVal_avgValDay_high_Day10'
**'dpData_meanVal_declinePT_duration'**
**'dpData_meanVal_relativeHighPeriod_duration'**
**'PCData_maxValDay_Day0'**
**'PCData_maxValDay_Day1'**
**'PCData_maxValDay_Day2'**
**'PCData_maxValDay_Day3'**
**'PCData_maxValDay_Day4'**
**'PCData_maxValDay_Day5'**
**'PCData_maxValDay_Day6'**
**'PCData_maxValDay_Day7'**
**'PCData_maxValDay_Day8'**
**'PCData_maxValDay_Day9'**
**'PCData_maxValDay_Day10'**
**'PCData_avgValDay_Day0'**
**'PCData_avgValDay_Day1'**
**'PCData_avgValDay_Day2'**
**'PCData_avgValDay_Day3'**
**'PCData_avgValDay_Day4'**
**'PCData_avgValDay_Day5'**
**'PCData_avgValDay_Day6'**
**'PCData_avgValDay_Day7'**
**'PCData_avgValDay_Day8'**
**'PCData_avgValDay_Day9'**
**'PCData_avgValDay_Day10'**
'PCData_maxValDay_high_Day0'

**'freqData_maxVal_maxValDay_high_Day4'**
**'freqData_maxVal_maxValDay_high_Day5'**
**'freqData_maxVal_maxValDay_high_Day6'**
**'freqData_maxVal_maxValDay_high_Day7'**
**'freqData_maxVal_maxValDay_high_Day8'**
**'freqData_maxVal_maxValDay_high_Day9'**
**'freqData_maxVal_maxValDay_high_Day10'**
**'freqData_maxVal_avgValDay_high_Day0'**
**'freqData_maxVal_avgValDay_high_Day1'**
**'freqData_maxVal_avgValDay_high_Day2'**
**'freqData_maxVal_avgValDay_high_Day3'**
**'freqData_maxVal_avgValDay_high_Day4'**
**'freqData_maxVal_avgValDay_high_Day5'**
**'freqData_maxVal_avgValDay_high_Day6'**
**'freqData_maxVal_avgValDay_high_Day7'**
**'freqData_maxVal_avgValDay_high_Day8'**
**'freqData_maxVal_avgValDay_high_Day9'**
**'freqData_maxVal_avgValDay_high_Day10'**
**'freqData_maxVal_declinePT_duration'**
**'freqData_maxVal_relativeHighPeriod_duration'**
**'freqData_numSeg_maxValDay_Day0'**
**'freqData_numSeg_maxValDay_Day1'**
**'freqData_numSeg_maxValDay_Day2'**
**'freqData_numSeg_maxValDay_Day3'**
**'freqData_numSeg_maxValDay_Day4'**
**'freqData_numSeg_maxValDay_Day5'**
**'freqData_numSeg_maxValDay_Day6'**
**'freqData_numSeg_maxValDay_Day7'**
**'freqData_numSeg_maxValDay_Day8'**
**'freqData_numSeg_maxValDay_Day9'**
**'freqData_numSeg_maxValDay_Day10'**
**'freqData_numSeg_avgValDay_Day0'**
**'freqData_numSeg_avgValDay_Day1'**
**'freqData_numSeg_avgValDay_Day2'**
**'freqData_numSeg_avgValDay_Day3'**
**'freqData_numSeg_avgValDay_Day4'**
**'freqData_numSeg_avgValDay_Day5'**
**'freqData_numSeg_avgValDay_Day6'**
**'freqData_numSeg_avgValDay_Day7'**
**'freqData_numSeg_avgValDay_Day8'**
**'freqData_numSeg_avgValDay_Day9'**
**'freqData_numSeg_avgValDay_Day10'**
**'freqData_numSeg_maxValDay_high_Day0'**

**'PCData_maxValDay_high_Day1'**
**'PCData_maxValDay_high_Day2'**
**'PCData_maxValDay_high_Day3'**
**'PCData_maxValDay_high_Day4'**
**'PCData_maxValDay_high_Day5'**
**'PCData_maxValDay_high_Day6'**
**'PCData_maxValDay_high_Day7'**
**'PCData_maxValDay_high_Day8'**
**'PCData_maxValDay_high_Day9'**
**'PCData_maxValDay_high_Day10'**
**'PCData_avgValDay_high_Day0'**
**'PCData_avgValDay_high_Day1'**
**'PCData_avgValDay_high_Day2'**
**'PCData_avgValDay_high_Day3'**
**'PCData_avgValDay_high_Day4'**
**'PCData_avgValDay_high_Day5'**
**'PCData_avgValDay_high_Day6'**
**'PCData_avgValDay_high_Day7'**
**'PCData_avgValDay_high_Day8'**
**'PCData_avgValDay_high_Day9'**
**'PCData_avgValDay_high_Day10'**
**'PCData_declinePT_duration'**
**'PCData_relativeHighPeriod_duration'**
**'freqData_numSeg_maxValDay_high_Day1'**
**'freqData_numSeg_maxValDay_high_Day2'**
**'freqData_numSeg_maxValDay_high_Day3'**
**'freqData_numSeg_maxValDay_high_Day4'**
**'freqData_numSeg_maxValDay_high_Day5'**
**'freqData_numSeg_maxValDay_high_Day6'**
**'freqData_numSeg_maxValDay_high_Day7'**
**'freqData_numSeg_maxValDay_high_Day8'**
**'freqData_numSeg_maxValDay_high_Day9'**
**'freqData_numSeg_maxValDay_high_Day10'**
**'freqData_numSeg_avgValDay_high_Day0'**
**'freqData_numSeg_avgValDay_high_Day1'**
**'freqData_numSeg_avgValDay_high_Day2'**
**'freqData_numSeg_avgValDay_high_Day3'**
**'freqData_numSeg_avgValDay_high_Day4'**
**'freqData_numSeg_avgValDay_high_Day5'**
**'freqData_numSeg_avgValDay_high_Day6'**
**'freqData_numSeg_avgValDay_high_Day7'**
**'freqData_numSeg_avgValDay_high_Day8'**
**'freqData_numSeg_avgValDay_high_Day9'**
**'freqData_numSeg_avgValDay_high_Day10'**
**'freqData_numSeg_declinePT_duration'**
**'freqData_numSeg_relativeHighPeriod_duration'**

Below is a table of the selected variables for the LASSO model for the wild-type population cultured at 25°C at a food level of $OD_{6005}$, along with the corresponding least-squares regression coefficient values.

| Behavioral Metric | LSR Coefficient |
|---|---|
| 'ampData_maxVal_avgValDay_Day0' | 0.040235 |
| 'ampData_maxVal_maxValDay_high_Day7' | -0.00035 |
| 'ampData_maxVal_avgValDay_high_Day6' | 0.011228 |
| 'ampData_maxVal_avgValDay_high_Day8' | 0.026329 |
| 'csData_maxVal_avgValDay_Day1' | 0.007468 |
| 'csData_maxVal_maxValDay_high_Day1' | 0.024184 |
| 'csData_maxVal_declinePT_duration' | 0.199274 |
| 'dpData_maxVal_relativeHighPeriod_duration' | -0.39457 |
| 'freqData_numSeg_avgValDay_high_Day8' | 0.027288 |
| 'PCData_avgValDay_Day8' | 0.047573 |
| 'PCData_avgValDay_Day9' | 0.053078 |
| 'PCData_declinePT_duration' | 3.66405 |
| 'PCData_relativeHighPeriod_duration' | -3.44935 |
